## Supplementary material for "SAMHD1 is a key regulator of the lineage-specific response of acute lymphoblastic leukaemias to nelarabine": Suppl Figure 1

### A

Heatmap illustrating expression (mRNA abundance) of all genes in B- vs. T-ALL cells based on CTRP data.

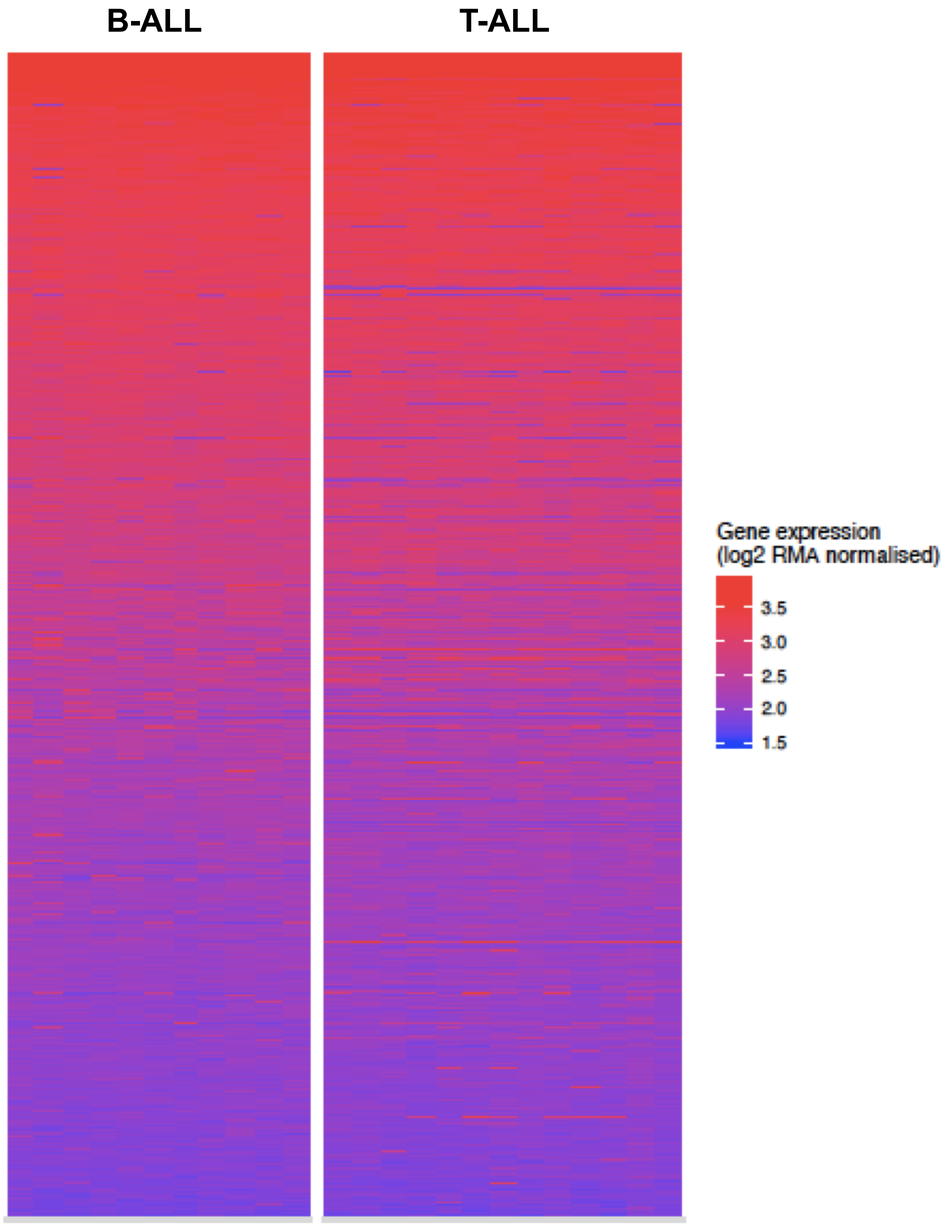

### Suppl. Figure 1

## B

Heatmap illustrating expression (mRNA abundance) of differentially regulated genes in B- vs. T-ALL cells based on CCLE data.

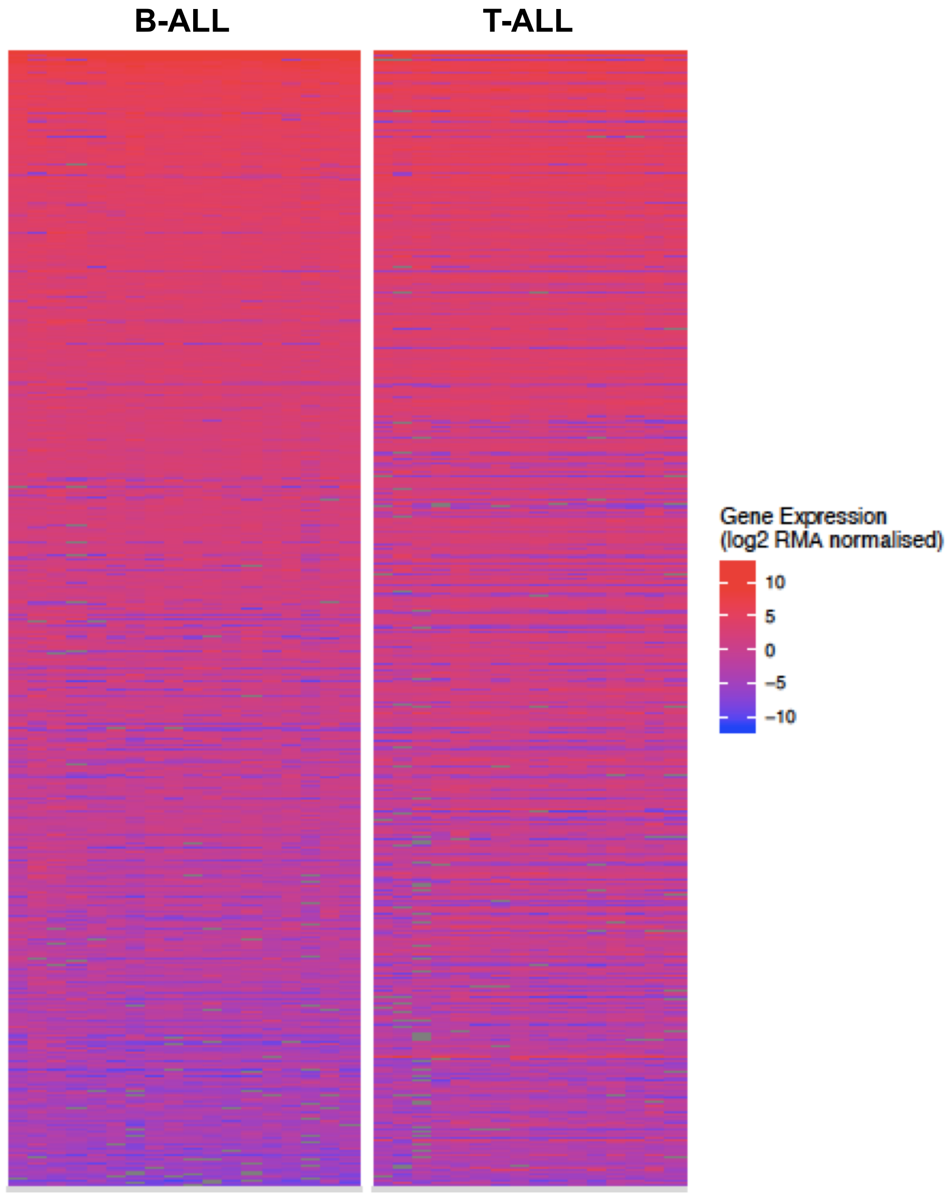

### Suppl. Figure 1

### C

Heatmap illustrating expression (mRNA abundance) of all genes in B- vs. T-ALL cells based on CCLE data.

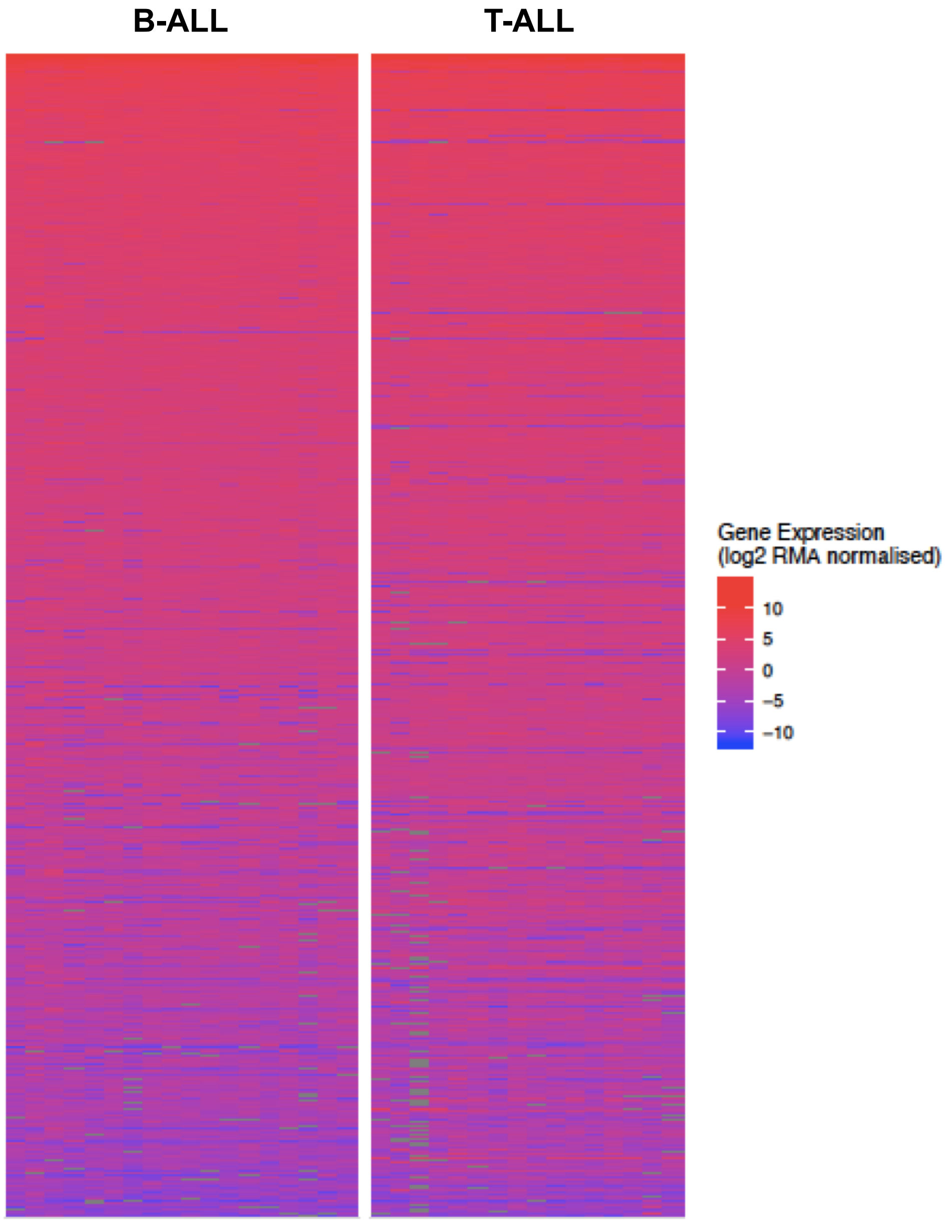

### Suppl. Figure 1

### D

Heatmap illustrating expression (mRNA abundance) of differentially regulated genes in B- vs. T-ALL cells based on GDSC data.

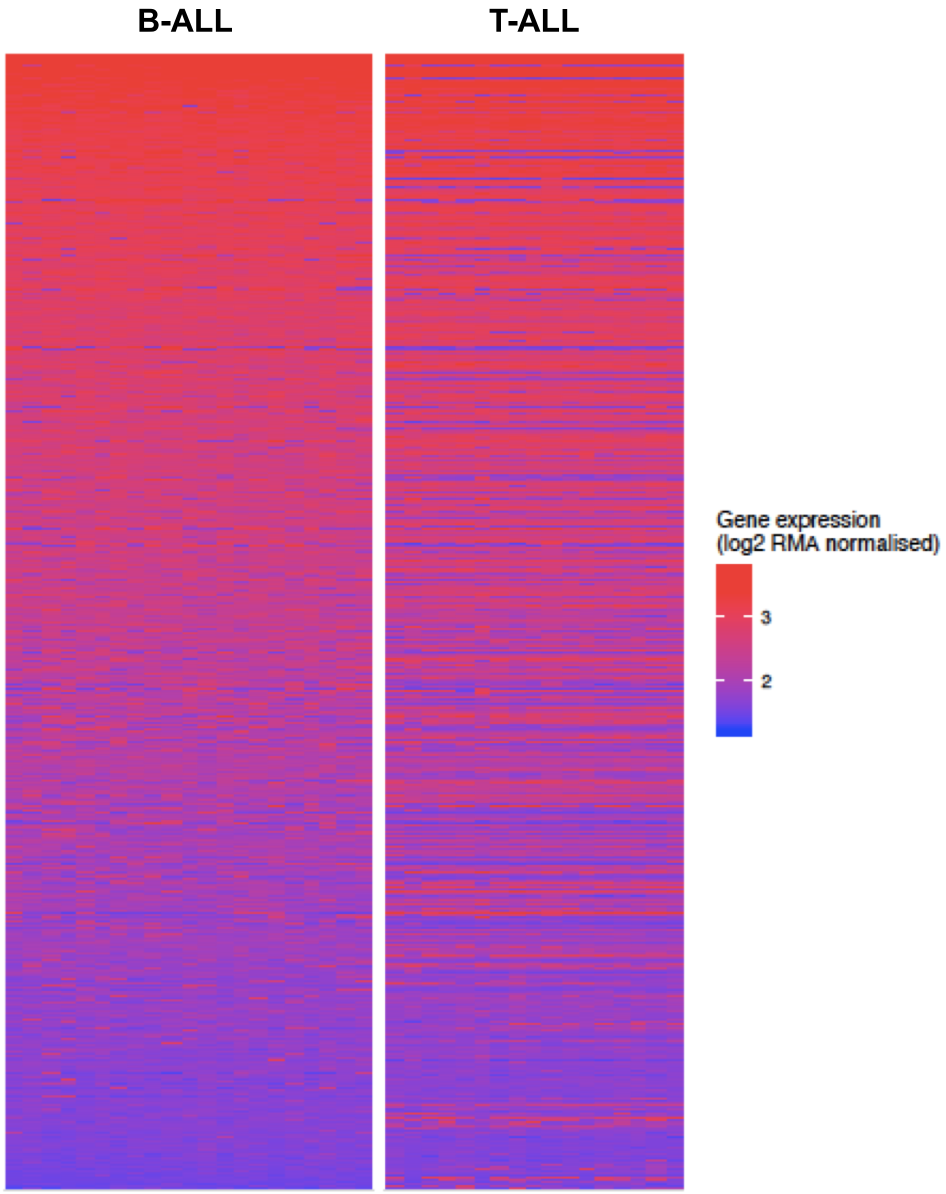

### Suppl. Figure 1

## E

Heatmap illustrating expression (mRNA abundance) of all genes in B- vs. T-ALL cells based on GDSC data.

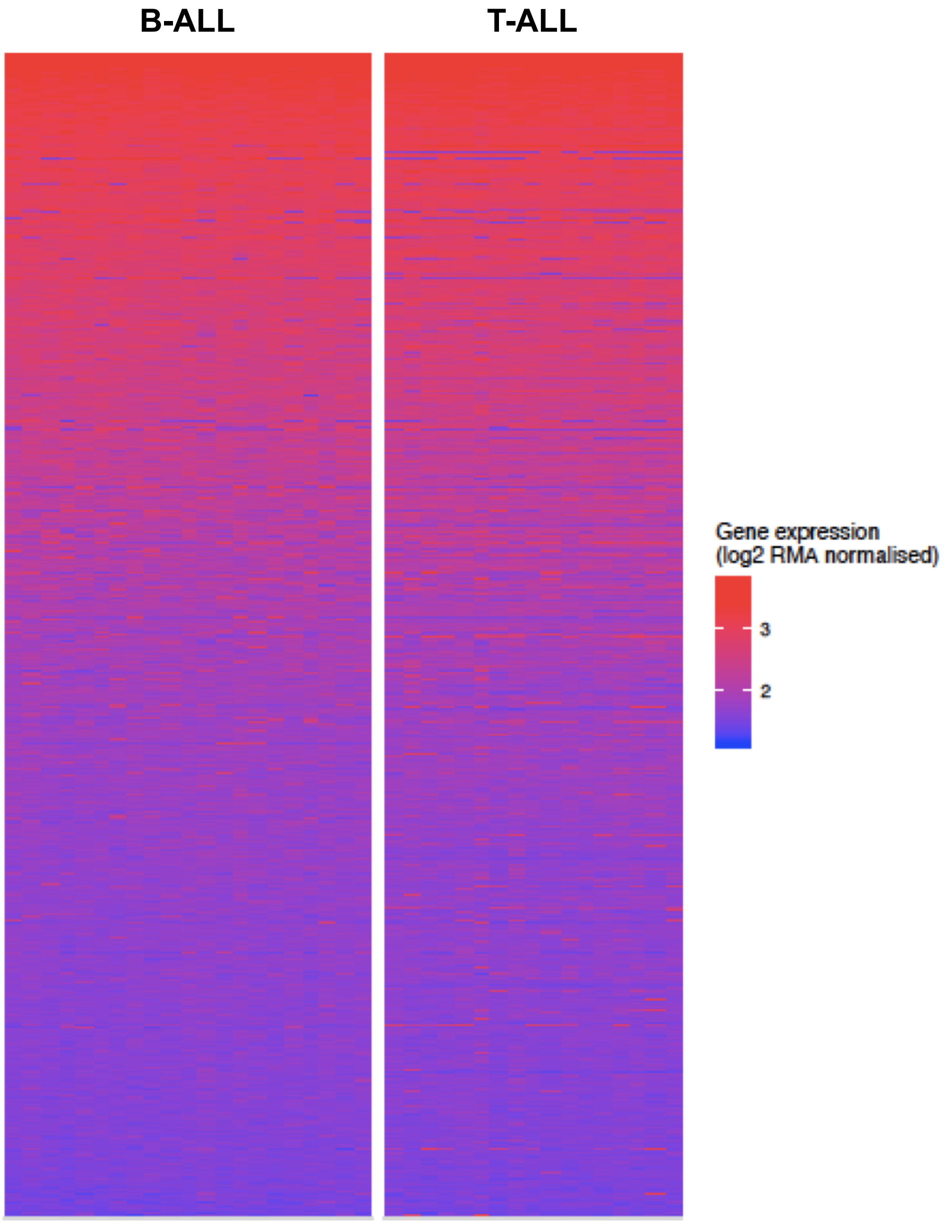
