## Supplementary figures and images for "SAMHD1 is a key regulator of the lineage-specific response of acute lymphoblastic leukaemias to nelarabine"

### Suppl Figure 3

**Suppl. Figure 3**

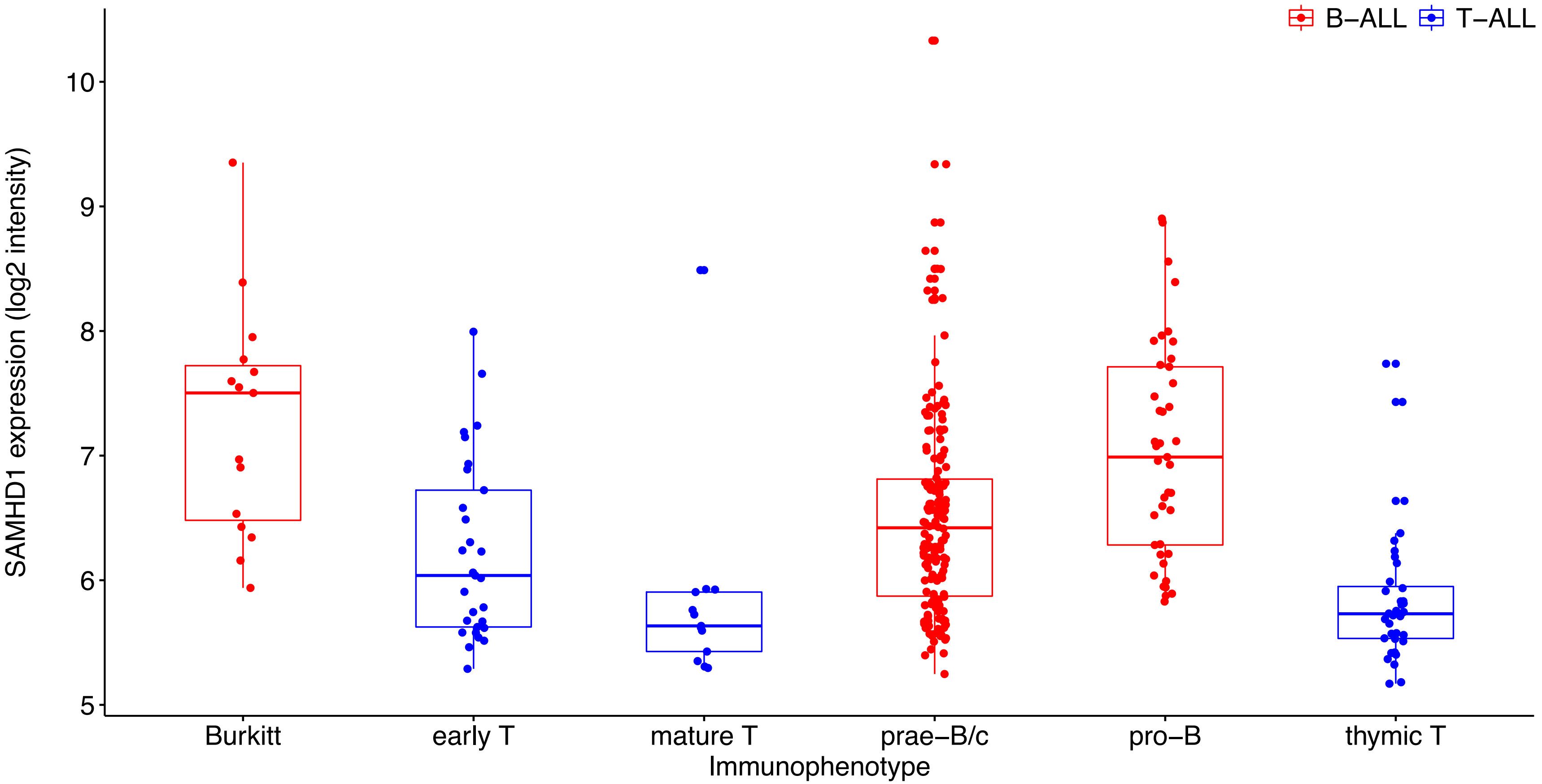


**Suppl. Figure 3**. SAMHD1 expression in ALL patients with different immunophenotypes.
