## Supplementary material for "SAMHD1 is a key regulator of the lineage-specific response of acute lymphoblastic leukaemias to nelarabine": Suppl Figure 4

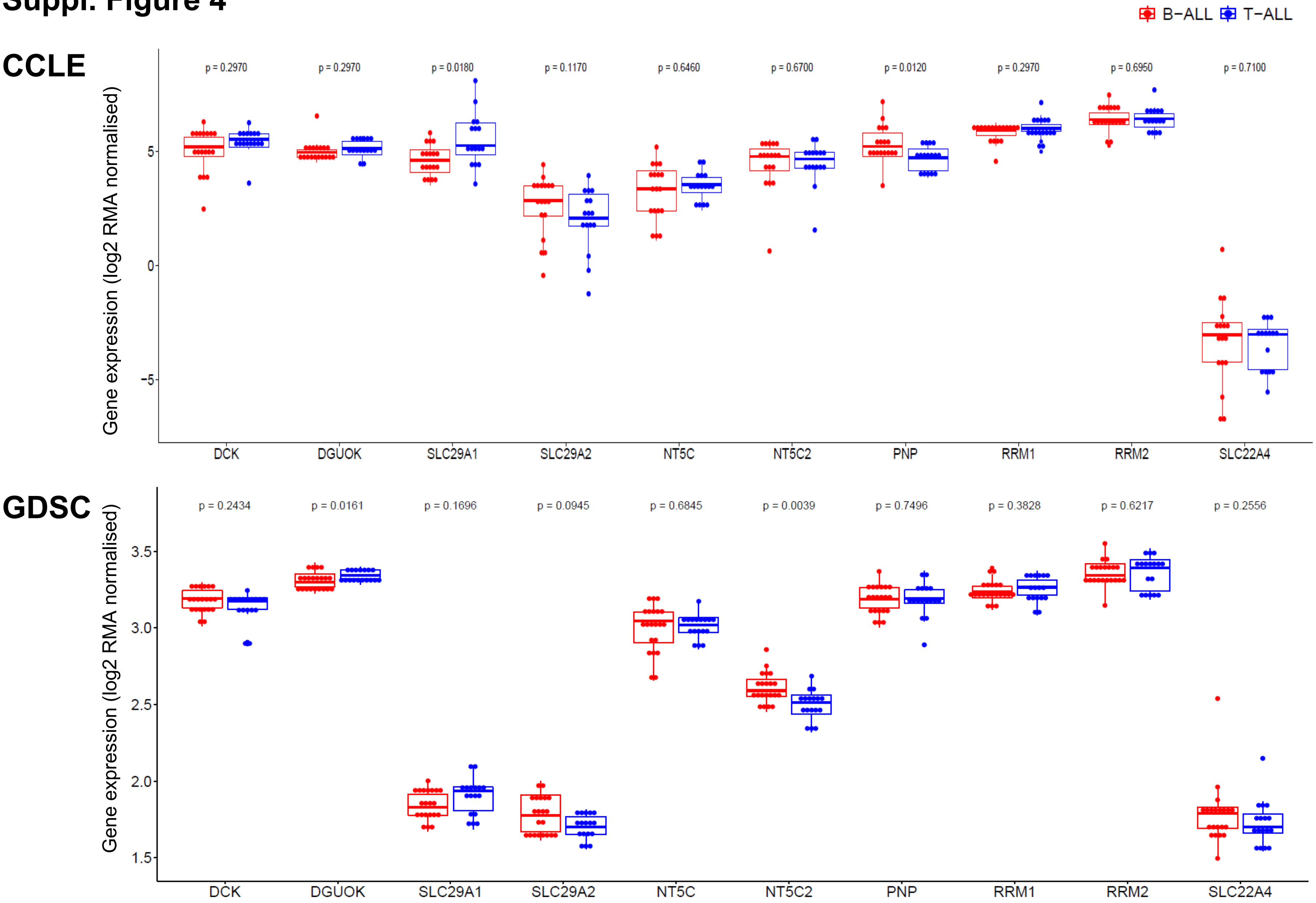


**Suppl. Figure 4**. Expression of genes known to be potentially involved in nucleoside analogue activity in B-ALL and T-ALL cell lines in the CCLE and GDSC.
