## Supplementary material for "SAMHD1 is a key regulator of the lineage-specific response of acute lymphoblastic leukaemias to nelarabine": Suppl Figure 5

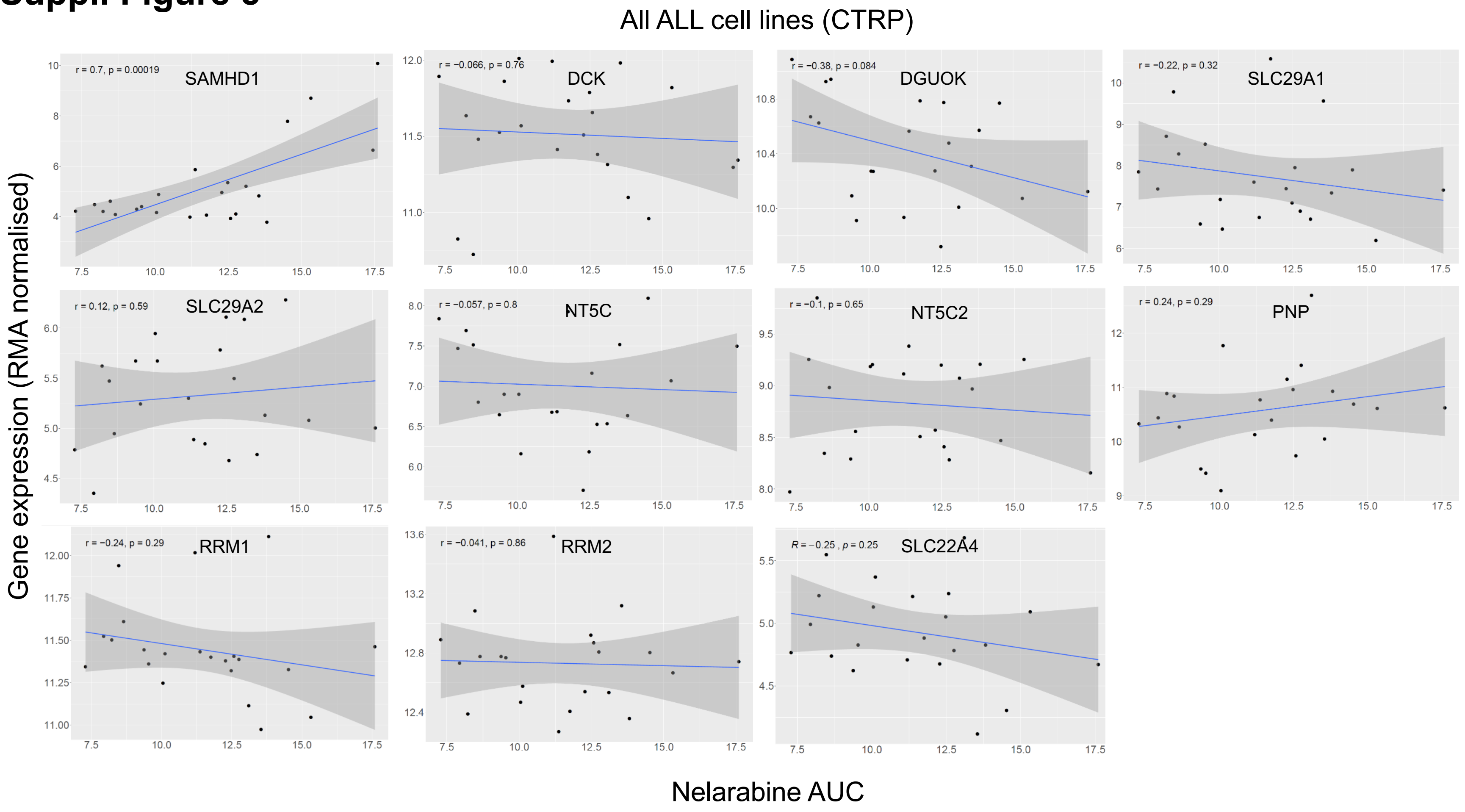

### Suppl. Figure 5

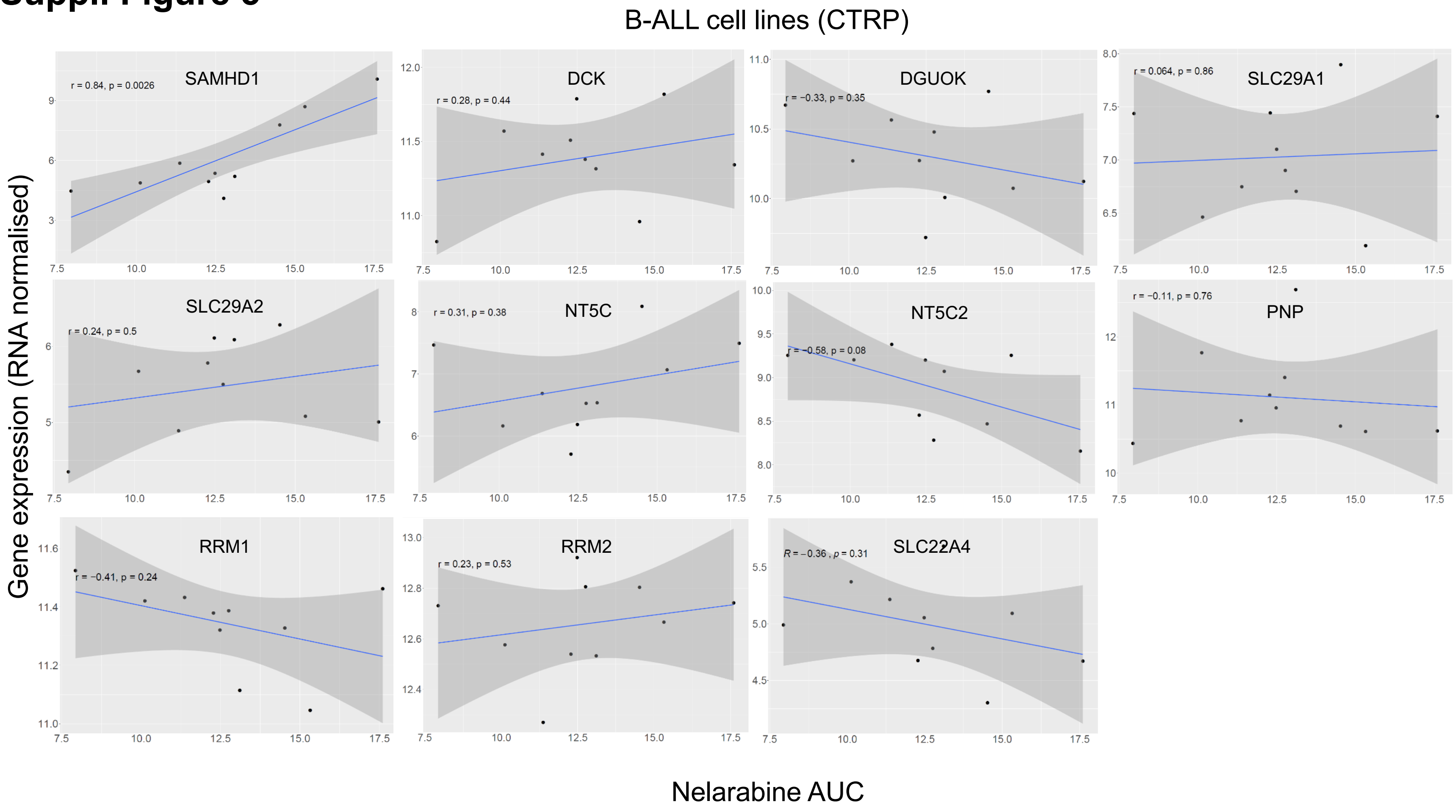

### Suppl. Figure 5

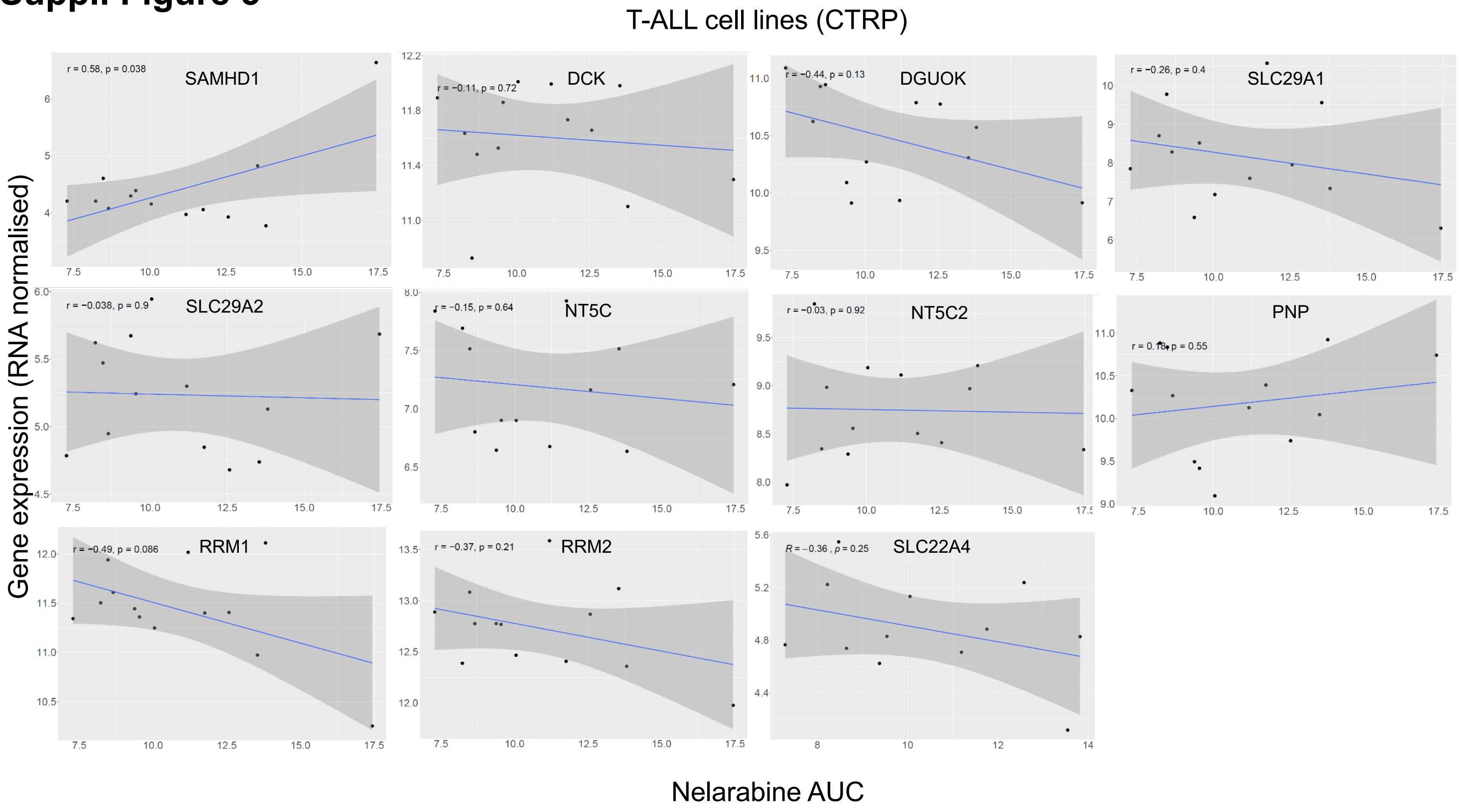


**Suppl. Figure 5**. Correlation of the expression of genes (mRNA abundance) known to affect nucleoside analogue activity to the nelarabine sensitivity (expressed as AUC) across all ALL, the B-ALL and the T-ALL cell lines based on CTRP data. Pearson's r values and respective p-values are provided.
