## Supplementary material for "SAMHD1 is a key regulator of the lineage-specific response of acute lymphoblastic leukaemias to nelarabine": Suppl Figure 6

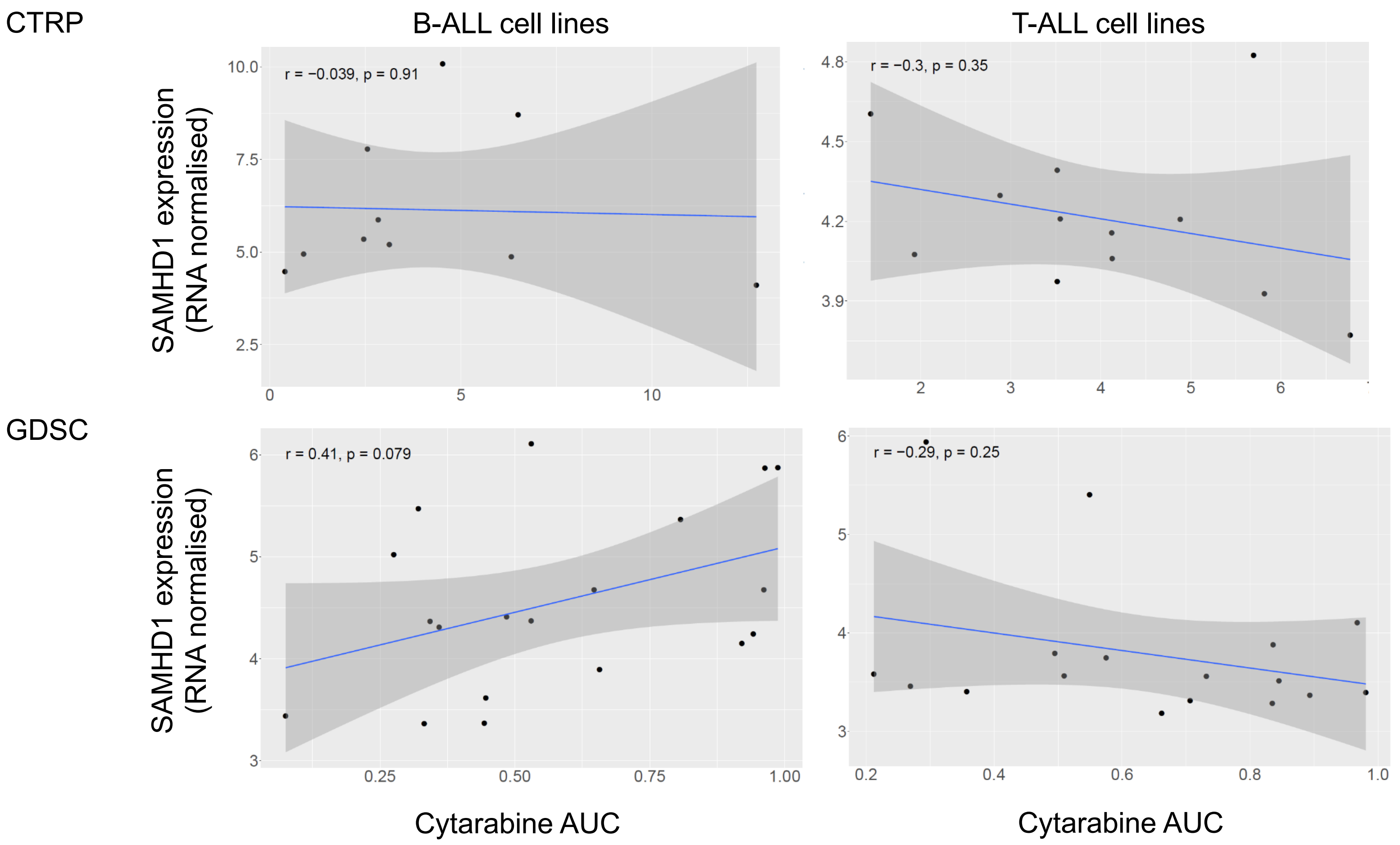


**Suppl. Figure 6**. Correlations of SAMHD1 expression (mRNA abundance) with the cytarabine AUC exclusively in B-ALL- and T-ALL cell lines based on CTRP and GDSC data. Pearson's r values and respective p-values are provided.
