## Supplementary material for "SAMHD1 is a key regulator of the lineage-specific response of acute lymphoblastic leukaemias to nelarabine": Suppl Figure 7

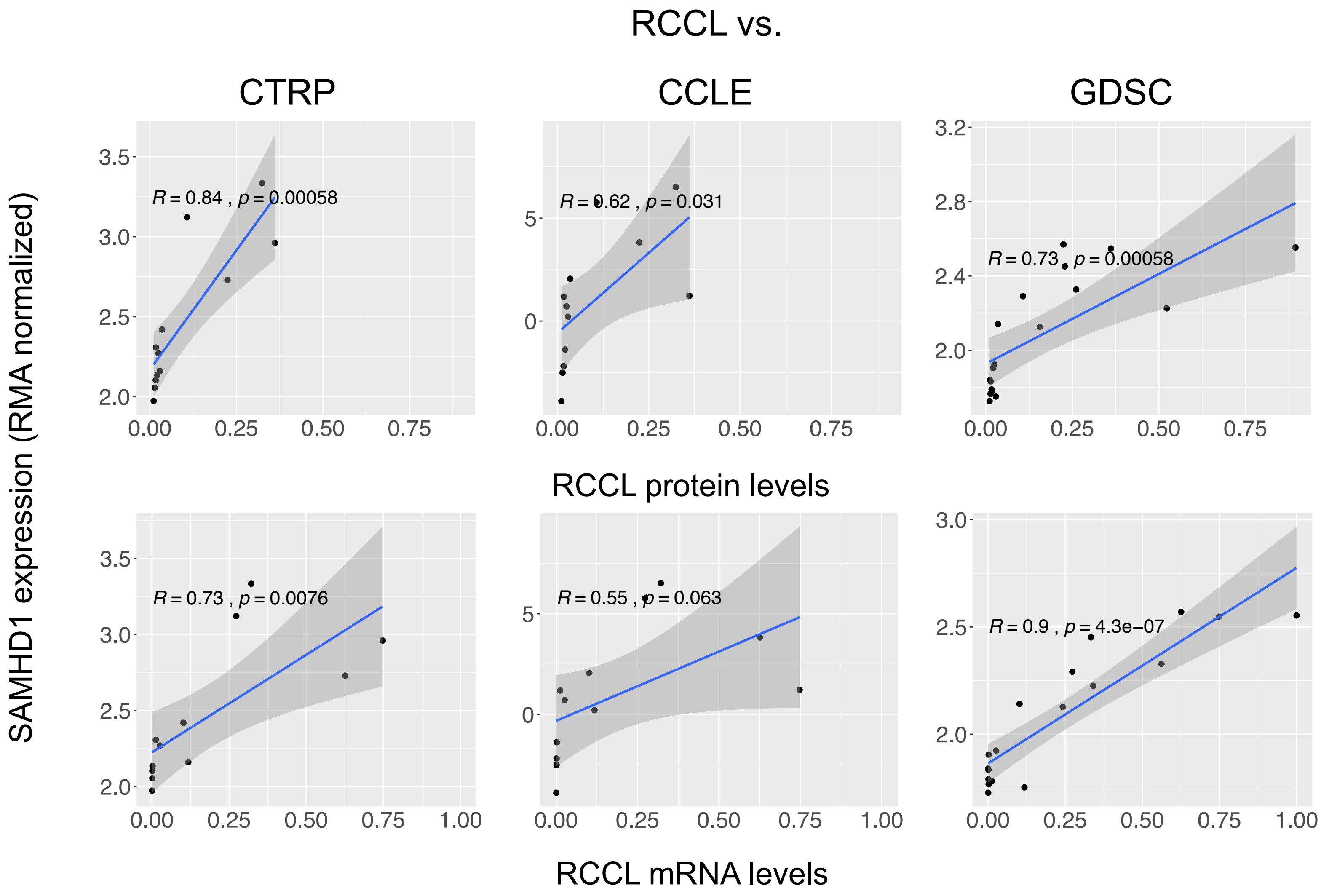


**Suppl. Figure 7**. Correlations of SAMHD1 protein and mRNA levels determined in the RCCL cell lines with the SAMHD1 expression data derived from the CTRP, CCLE, and GDSC among the cell lines that are represented in both respective datasets. Pearson's r values and respective p-values are provided.
