## Supplementary material for "SAMHD1 is a key regulator of the lineage-specific response of acute lymphoblastic leukaemias to nelarabine": Suppl Figure 8

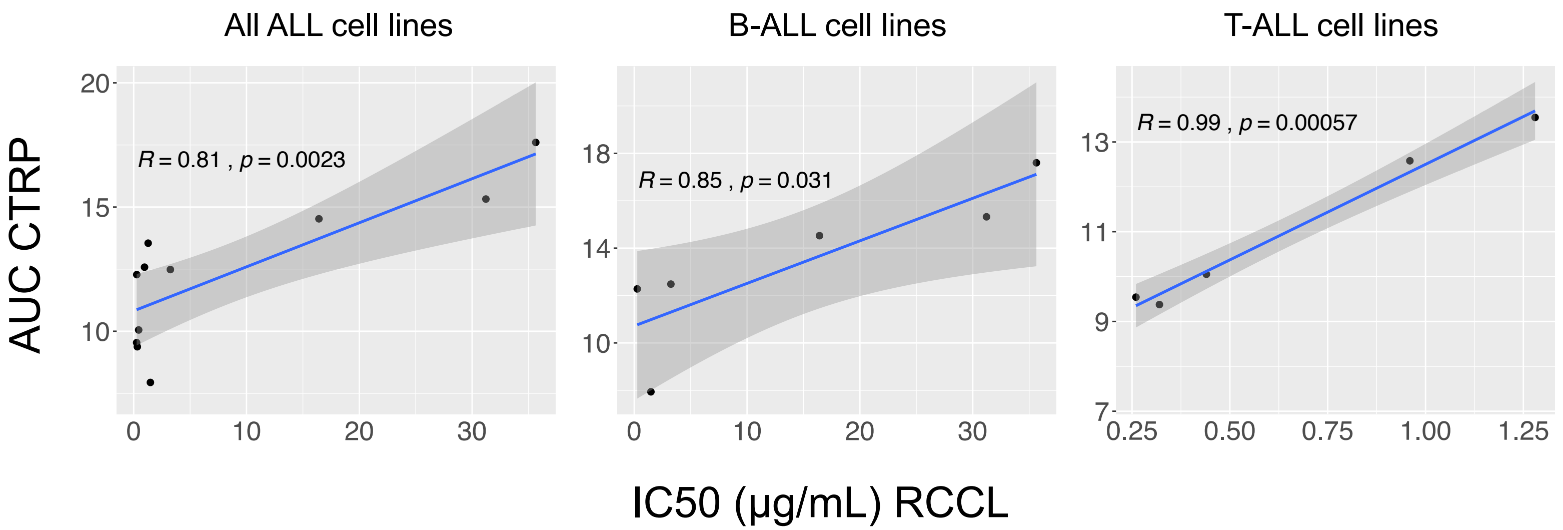


**Suppl. Figure 8**. Correlations of the nelarabine AUCs derived from the CTRP and the AraG IC50 values determined in the RCCL panel across the ALL cell lines present in both datasets. Pearson's r values and respective p-values are provided.
