## Supplementary material for "SAMHD1 is a key regulator of the lineage-specific response of acute lymphoblastic leukaemias to nelarabine": Suppl Table 1

**Suppl. Table 1**. B-ALL and T-ALL cell lines in the CCLE and GDSC (overlaps highlighted in *italics*).

| **CCLE** | | **GDSC** | |
|---|---|---|---|
| **Cell Line** | **Lineage** | **Cell line** | **Lineage** |
| *697* | B-ALL | *697* | B-ALL |
| A4FUK | B-ALL | ALL-PO | B-ALL |
| EHEB | B-ALL | BALL-1 | B-ALL |
| HUNS1 | B-ALL | GR-ST | B-ALL |
| JM-1 | B-ALL | HAL-01 | B-ALL |
| KASUMI-2 | B-ALL | KARPAS-231 | B-ALL |
| *KOPN-8* | B-ALL | *KOPN-8* | B-ALL |
| *MHH-CALL2* | B-ALL | LC4-1 | B-ALL |
| MHH-CALL3 | B-ALL | *MHH-CALL-2* | B-ALL |
| *MHH-CALL4* | B-ALL | *MHH-CALL-4* | B-ALL |
| MUTZ-5 | B-ALL | MHH-PREB-1 | B-ALL |
| NALM-19 | B-ALL | MN-60 | B-ALL |
| *NALM-6* | B-ALL | *NALM-6* | B-ALL |
| *RCH-ACV* | B-ALL | P30-OHK | B-ALL |
| *REH* | B-ALL | *RCH-ACV* | B-ALL |
| *RS-411* | B-ALL | *REH* | B-ALL |
| SEM | B-ALL | ROS-50 | B-ALL |
| *SUP-B15* | B-ALL | *RS4-11* | B-ALL |
| | | *SUP-B15* | B-ALL |
| | | SUP-B8 | B-ALL |
| | | U-698-M | B-ALL |
| *ALL-SIL* | T-ALL | *ALL-SIL* | T-ALL |
| C8166 | T-ALL | ATN-1 | T-ALL |
| *DND-41* | T-ALL | BE-13 | T-ALL |
| HPB-ALL | T-ALL | CCRF-CEM | T-ALL |
| *JURKAT* | T-ALL | *DND-41* | T-ALL |
| *KE-37* | T-ALL | HH | T-ALL |
| *LOUCY* | T-ALL | *JURKAT* | T-ALL |
| *MOLT-13* | T-ALL | KARPAS-45 | T-ALL |
| *MOLT-16* | T-ALL | *KE-37* | T-ALL |
| MOLT-3 | T-ALL | *LOUCY* | T-ALL |
| *P12-ICHIKAWA* | T-ALL | *MOLT-13* | T-ALL |
| PEER | T-ALL | *MOLT-16* | T-ALL |
| *PF-382* | T-ALL | MOLT-4 | T-ALL |
| *RPMI-8402* | T-ALL | *P12-ICHIKAWA* | T-ALL |
| *SUP-T11* | T-ALL | *PF-382* | T-ALL |
| TALL-1 | T-ALL | *RPMI-8402* | T-ALL |
| | | *SUP-T11* | T-ALL |
