## Supplementary material for "SAMHD1 is a key regulator of the lineage-specific response of acute lymphoblastic leukaemias to nelarabine": Suppl Table 2

**Suppl. Table 2**. B-ALL and T-ALL cell line sensitivity to nelarabine expressed as area under the curve (AUC) derived from CTRP.

| Cell line | Lineage | AUC |
|---|---|---|
| HPBALL | T-ALL | 11.757 |
| DND-41 | T-ALL | 7.29 |
| SUPT-1 | T-ALL | 8.4742 |
| JURKAT | T-ALL | 12.58 |
| PEER | T-ALL | 13.818 |
| PF-382 | T-ALL | 11.193 |
| ALL-SIL | T-ALL | 13.546 |
| P12-ICHIKAWA | T-ALL | 9.5418 |
| RPMI-8402 | T-ALL | 10.052 |
| MOLT-16 | T-ALL | 17.44 |
| MOLT-13 | T-ALL | 8.6473 |
| TALL-1 | T-ALL | 8.2253 |
| KE-37 | T-ALL | 9.3765 |
| SEM | B-ALL | 17.602 |
| RCH-ACV | B-ALL | 12.759 |
| MHH-CALL3 | B-ALL | 10.123 |
| RS-411 | B-ALL | 14.526 |
| MHH-CALL4 | B-ALL | 15.322 |
| REH | B-ALL | 7.9391 |
| 697 | B-ALL | 12.28 |
| SUP-B15 | B-ALL | 11.372 |
| KASUMI-2 | B-ALL | 13.107 |
| NALM-6 | B-ALL | 12.483 |
