## Supplementary material for "SAMHD1 is a key regulator of the lineage-specific response of acute lymphoblastic leukaemias to nelarabine": Suppl Table 6

**Suppl. Table 6**. AraG and cytarabine concentrations that reduce B-ALL and T-ALL cell line sensitivity by 50% (IC50).

| **B-ALL** | **IC50** | |
|---|---|---|
| **Cell line** | **AraG (µg/mL)** | **Cytarabine (ng/mL)** |
| 697 | 0.27 ± 0.01 | 1.23 ± 0.05 |
| BALL-1 | 4.76 ± 0.50 | 2.74 ± 0.05 |
| GRANTA-452 | 6.69 ± 0.70 | 3.42 ± 0.25 |
| HAL-01 | 2.15 ± 0.49 | 0.98 ± 0.04 |
| KARPAS231 | 26.73 ± 2.62 | 11.57 ± 0.77 |
| MHH-CALL-4 | 31.22 ± 2.50 | 27.07 ± 3.05 |
| MN-60 | 99.10 ± 1.62 | 14.95 ± 0.99 |
| NALM-6 | 3.25 ± 0.28 | 2.19 ± 0.06 |
| NALM-16 | 65.19 ± 2.72 | 8.79 ± 0.42 |
| REH | 1.48 ± 0.07 | 1.31 ± 0.17 |
| ROS-50 | 90.62 ± 9.05 | 18.28 ± 3.57 |
| RS4;11 | 16.42 ± 1.32 | 5.74 ± 1.89 |
| SEM | 35.64 ± 3.71 | 27.35 ± 3.18 |
| TANOUE | 49.14 ± 2.95 | 27.67 ± 0.60 |
| TOM-1 | 0.10 ± 0.01 | 1.38 ± 0.03 |
| **T-ALL** | **IC50** | |
| **Cell line** | **AraG (µg/mL)** | **Cytarabine (ng/mL)** |
| ALL-SIL | 1.28 ± 0.16 | 5.17 ± 0.64 |
| CCRF-CEM | 0.43 ± 0.02 | 3.88 ± 0.70 |
| CTV-1 | 0.38 ± 0.07 | 1.72 ± 0.01 |
| HSB-2 | 0.52 ± 0.05 | 3.51 ± 0.07 |
| JJHan | 0.97 ± 0.19 | 5.61 ± 0.53 |
| Jurkat | 0.96 ± 0.06 | 6.65 ± 0.52 |
| KE-37 | 0.32 ± 0.11 | 1.83 ± 0.25 |
| MOLT-4 | 0.46 ± 0.01 | 3.02 ± 0.10 |
| MOLT-16 | 15.55 ± 1.30 | 6.48 ± 0.53 |
| P12-ICHIKAWA | 0.26 ± 0.01 | 2.12 ± 0.19 |
| RPMI-8402 | 0.44 ± 0.01 | 3.43 ± 0.21 |
